# NOX2 Inhibition Enables Retention of the Circadian Clock in BV2 Microglia and Primary Macrophages

**DOI:** 10.1101/2022.11.07.515487

**Authors:** Iswarya Muthukumarasamy, Sharleen M. Buel, Jennifer M. Hurley, Jonathan S. Dordick

## Abstract

Sustained neuroinflammation is a major contributor to the progression of neurodegenerative diseases such as Alzheimer’s (AD) and Parkinson’s (PD) diseases. Neuroinflammation, like other cellular processes, is affected by the circadian clock. Microglia, the resident immune cells in the brain, act as major contributors to neuroinflammation and are under the influence of the circadian clock. Microglial responses such as activation, recruitment, and cytokine expression are rhythmic in their response to various stimuli. While the link between circadian rhythms and neuroinflammation is clear, significant gaps remain in our understanding of this complex relationship. To further our understanding of this relationship, we studied the interaction between the microglial circadian clock and the enzyme NADPH Oxidase Isoform 2 (NOX2), an enzyme essential for the production of reactive oxygen species (ROS) in oxidative stress, an integral characteristic of neuroinflammation. We examined BV2 microglia over circadian time, demonstrating oscillations of the clock genes *Per2* and *Bmal1* and the NOX2 subunits *gp91^phox^* and *p47^phox^*. We discovered the BV2 microglial clock exerted significant control over NOX2 expression and that the inhibition of NOX2 enabled the microglia to retain a functional circadian clock while reducing levels of ROS and inflammatory cytokines. These trends were mirrored in mouse bone marrow-derived primary macrophages. Our findings indicate NOX2 plays a crucial role in the interaction between the circadian clock and the activation of microglia/macrophages into their pro-inflammatory state.

## INTRODUCTION

Microglia, the innate immune cells of the central nervous system, are involved in inflammation and the immune response, and play a significant role in contributing to neuroinflammation associated with neurodegenerative diseases (1). Like macrophages, microglia display multiple phenotypes based on external stimuli, and are normally distinguished into resting and activated states in the brain (2). In their resting state, microglia carry out continuous surveillance of the central nervous system to maintain homeostasis (2). The activated microglial state is itself an oversimplification and comprises multiple phenotypes (3,4), broadly differentiated into pro-inflammatory and anti-inflammatory states (3,4). The pro-inflammatory phenotype is associated with the release of mediators that lead to inflammation (5–7), whereas the anti-inflammatory phenotype is associated with the release of mediators that promote wound healing and debris clearance (4,8,9). Under normal healthy conditions, there is a fine balance between these phenotypes, which ensures homeostasis. However, in the case of various neurodegenerative diseases, the intricate balance among microglial phenotypes is lost and there is sustained activation of microglia into a pro-inflammatory state, resulting in neuroinflammation (1,4). Such chronic neuroinflammation has been associated with the progression of various neurodegenerative diseases such as Alzheimer’s (AD) and Parkinson’s (PD) diseases, among others, making neuroinflammation a potential target for therapeutic intervention.

A predominant aspect of the microglial pro-inflammatory response is the release of reactive oxygen species (ROS), which is facilitated by the enzyme NADPH oxidase isoform 2 (NOX2) (10). ROS generated by NOX2 is normally under tight control of various protective enzymes, including catalases, superoxide dismutases, and glutathione peroxidases, as well as small molecule antioxidants (11). When such regulatory processes fail, the cell enters a state of oxidative stress (12). The overproduction of ROS combined with the reduced activity of ROS scavenging enzymes results in oxidative damage of biomolecules and neurons (11,13,14), making NOX2 an attractive target to combat chronic neuroinflammation (15,16).

The immune responses, including neuroinflammatory responses, are tightly controlled by the circadian clock (17). Circadian clocks are broadly present in organisms that live in the photic zone and ensure efficient synchronization between physiologic functions and the 24 h structure of the day (18). This time-based regulation by the circadian clock is facilitated through a transcriptional-translational feedback loop (TTFL) orchestrated by a pair of transcriptional activators/repressors. Through this intricate feedback loop, the circadian clock ensures the oscillation of a broad array of genes and proteins to time cellular physiology. At the cellular level, macrophages and monocytes have robust circadian clocks with high amplitude oscillations of their core clock genes and proteins (19,20). This strong influence of the circadian clock over immune cell physiology results in 24 h rhythms in levels of cytokines, recruitment of macrophages and monocytes to tissues, phagocytosis, and response of pattern recognition receptors (17). Paralleling what is observed in macrophages, microglia also display high amplitude rhythms in circadian clock genes and inflammatory cytokines (21,22).

Disruption of the circadian clock has been associated with increased risk for, and progression of, disease states, particularly diseases with inflammatory components (18,23–26), including neurodegenerative diseases (26,27). Mouse models have shown that impairment of the microglial clock results in increased levels of inflammation and worsening of AD symptoms (28). In parallel, activation of macrophages into pro- and anti-inflammatory phenotypes directly modulates the function of the circadian clock (29), indicating the complex relationship between inflammatory responses, the circadian clock, and disease states. Although the fields of circadian biology and neuroinflammation have advanced exponentially in recent years, there remain significant gaps in our understanding of the interrelationship between circadian rhythms and neuroinflammation. Gaining a mechanistic insight into this intricate relationship may advance the development of interventional therapies that mitigate circadian disruption thereby preventing neuroinflammation associated with various neurodegenerative diseases.

The strong dependence of neuroinflammation on NOX2 activation (13,30–32) and circadian disruption (17,19,33,34) has been studied individually, but the correlation between NOX2 activation and circadian disruption has yet to be explored in depth. In the current study, the influence of the circadian clock in a BV2 microglial cell line has been investigated, specifically with respect to NOX2 expression. Activation of BV2 microglia into pro- or anti-inflammatory phenotypes resulted in distinct effects similar to those observed in macrophages, with the clock losing its rhythm under pro-inflammatory activation and retaining its rhythm under anti-inflammatory activation (29). Furthermore, the circadian clock in BV2 microglia exerted strong control over the expression of various NOX2 enzyme subunits. Inhibition of NOX2 in a pro-inflammatory state resulted in retention of a functional circadian clock, suggesting that NOX2 plays an important role in the connection between neuroinflammation and circadian disruption in the context of neurodegenerative diseases. Finally, we found the link between the clock and NOX2 in BV2 cells consistent with data from primary bone marrow-derived mouse macrophages, indicating a conserved connection between NOX2 and the clock in cells involved in the inflammatory response.

## MATERIALS AND METHODS

### Materials

The BV2 mouse microglial cell line was obtained from Banca Cellule ICLC, Genova, Italy. Lipopolysaccharide (LPS) 500X was obtained from Invitrogen (Waltham, MA). Murine IL-4 was obtained from Peprotech Inc. (Rocky Hill, NJ). NOX2 inhibitors apocynin and GSK2795039 (GSK) were purchased from Millipore Sigma (Burlington, MA) and MedChemExpress (Monmouth Junction, NJ), respectively. Primary antibodies against phospho-p47^*phox*^ (Ser 370), PER2, and BMAL1, and goat anti-rabbit secondary antibody were obtained from Invitrogen. Phosphate buffered saline (PBS) and trypsin-EDTA for cell culture were obtained from Gibco, ThermoFisher Scientific (Waltham, MA). The Amido Black stain was obtained from ThermoFisher Scientific.

### BV2 Cell Culture and Synchronization

BV2 cells were cultured in RPMI-1640 medium (ThermoFisher Scientific) with L-glutamine (2 mM) and supplemented with 10% heat-inactivated FBS (ThermoFisher Scientific). All circadian studies used a serum-shock method to synchronize the cells (20). The cells were plated in normal growth media (RPMI + 10% FBS) at a density of 10^5^ cells/mL in 6-well plates and allowed to reach 90% confluence. Culture media was then replaced with starvation media (RPMI) for 24 h. Following media starvation, the cells were subjected to serum-shock using 50% FBS for 2 h to synchronize their clock (20,35–37). Post-serum shock, normal growth media was used, and the cells were allowed to recover for 16 h from the serum-shock and attain homeostasis. Samples were collected every 2 h for 24 h, starting at 16 h post serum-shock (Hours Post Serum Shock 16 or HPS16), and processed for RT-qPCR and Western blotting.

### Bone marrow derived macrophage (BMDMs) extraction and synchronization

Bone marrow was extracted from the tibias and femurs of 3-6 month old male *Per2:Luc* (C57BL/6J) mice. The bone marrow progenitor cells were differentiated with DMEM supplemented with M-CSF and 10%FBS as mentioned in a previous protocol (20,37). After 7 days, the differentiated macrophages were synchronized by a 24 h starve in serum-free media followed by a 2 h serum-shock with 50% FBS as mentioned previously (20,37). Luminescence was measured using LumiCycle32 (Actimetrics, Wilmette, IL) with cells plated in Leibovitz media containing Luciferin and 10% FBS (20,37).

### Quantitative real-time PCR

The High Pure RNA Isolation Kit from Roche Molecular Systems, Inc. (Branchburg, NJ) was used for RNA extraction and purification. Purified RNA was reverse-transcribed using the High-Capacity cDNA Reverse Transcription Kit (ThermoFisher Scientific). mRNA abundance was determined by quantitative PCR using TaqMan Gene Expression Master Mix (Applied Biosystems, Waltham, MA) and the following pre-designed TaqMan primer assays: *Per2* (Assay ID: Mm00478099_m1), *Bmal1* (Assay ID: Mm00500223_m1), *Hprt1* (Assay ID: Mm03024075_m1), *p47^phox^* (Assay ID: Mm00447921_m1) and *gp91^phox^* (Assay ID: Mm01287743_m1). *Hprt1* was used as the reference gene for all RT-qPCR experiments. Data obtained from RT-qPCR was then input into the ECHO software (see Supplementary Information) for analysis of oscillation of gene expression (38).

### Western blotting

Western blotting was used to investigate the oscillation of PER2 and BMAL1 clock proteins, and for quantification of phosphorylated-p47^*phox*^ levels in BV2 microglia. Time course samples were collected from synchronized BV2 cells as described above. For measurement of phospho-p47^*phox*^ levels, BV2 cells were plated in 6-well plates at a density of 10^5^ cells/mL and allowed to reach 90% confluence. At confluence, cells were treated with LPS (1 μg/mL), or LPS (1 μg/mL) + apocynin (100 μM), or LPS (1 μg/mL) + GSK2795039 (25 μM) for 24 h. BV2 cells with no additives were used as a control. For all samples, cells were washed with ice-cold PBS and lysed with RIPA lysis buffer (Millipore Sigma, Burlington, MA) containing the Halt Protease Inhibitor Cocktail (ThermoFischer Scientific). Protein concentrations were measured using the Pierce BCA Protein Assay Kit (Pierce, Waltham, MA). Samples were loaded at 25 μg of total protein per lane, resolved by SDS-PAGE (BioRad, Hercules, CA), and transferred onto nitrocellulose membranes (Bio-Rad). Following blocking with 5% skim milk in PBST (137 mM NaCl, 2.7 mM KCl, 10 mM Na_2_HPO_4_, 1.8 mM KH_2_PO_4_, pH 7.4, 0.2% Tween-20) for 2 h at room temperature, the membranes were incubated with primary antibodies against PER2 (1:1000, Invitrogen, Waltham, MA), BMAL1 (1:1000, Invitrogen), and phospho-p47^*phox*^ (Ser 370) (1:1000, Invitrogen) at 4°C in 1% skim milk in PBST overnight. Post-overnight incubation, the membranes were washed six times with PBST and incubated with secondary antibody (1:10,000) for 1 h. The membranes were then washed with PBST, and the protein bands were detected using SuperSignal West Pico Chemiluminescent Substrate (Pierce). Amido black staining (see below) was used to normalize protein loading. Protein bands were detected using the BioRad ChemiDoc XRS+ Imager (BioRad). Quantification of detected protein bands was performed with the ImageLab 6.0.1 software (Life Science, Waltham, MA).

### Amido Black staining

Amido Black staining was used as the internal loading-control for Western blots. Following detection of protein bands, membranes were washed three times in DI water. The membranes were then stained for 1 min using a staining solution containing 0.1% amido black reagent in 10% acetic acid solution. The membranes were then washed twice for 1 min with a de-staining solution containing 5% acetic acid. Post de-staining, the membranes were washed two times for 10 min each in DI water. The membranes were then air-dried and visualized using a Bio-Rad ChemiDoc XRS+ Imager. Quantification of amido black stained membranes was done using the ImageLab 6.0.1 software.

### ECHO analysis

ECHO version 4.1 was used to analyze all transcriptional and protein data obtained from the time-course experiments (38). All data was free run with the ‘smooth data’ and ‘linear detrend’ options selected. For transcriptional data, the ‘normalize data’ option was also selected. Data obtained from ECHO analysis was then transferred and re-plotted on GraphPad Prism 7 (GraphPad Software, CA, USA). Details about the ECHO software and ECHO data for all transcriptome and proteome oscillations are available in the Additional File 6: Table S1.

### ROS determination

ROS released by BV2 in the presence of LPS (1 μg/mL) and NOX2 inhibitors apocynin (100 μM) and GSK2795039 (25 μM) was measured using two different assays. The DCFDA/H2DCFDA Cellular ROS Assay Kit (Abcam, MA) was used to measure the levels of intracellular superoxide produced and the Amplex Red Hydrogen Peroxide/Peroxidase Assay kit (Invitrogen) was used to measure the levels of extracellular hydrogen peroxide produced in BV2 cells. Cells were plated in 96-well black-walled flat-bottom plates at a density of 10^5^ cells/mL. The cells were incubated with LPS with and without a NOX2 inhibitor for 2 h in the assay media and fluorescence was measured using a SpectraMax plate reader (Molecular Devices, San Jose, CA). Fluorescence values were normalized to cell number measured using the Presto Blue Cell Viability Assay (ThermoFischer Scientific) as described below.

### Cell viability determination

BV2 cells were seeded in 96-well flat-bottom, black-walled plates at a density of 10^4^ cells/well and allowed to reach 90% confluence. The cells were then treated with media containing different additives. The media was then removed from each well and replaced with 90 μL fresh media and 10 μL PrestoBlue reagent (ThermoFisher Scientific) and incubated at 37°C and 5% CO_2_ for 10 min. Fluorescence was measured at Ex/Em = 560/590 nm and cell numbers were correlated against a standard curve.

### ELISA

A sandwich enzyme-linked immunosorbent assay (ELISA) was used to quantify cytokines produced by BV2 cells and primary macrophages. The cells were plated in 96-well plates at a density of 10^5^ cells/mL and allowed to reach 90% confluence. The cells were then incubated for 24 h with media containing the following combination of additives: LPS (1 μg/mL); LPS (1 μg/mL) + apocynin (100 μM); apocynin (100 μM); LPS (1 μg/mL) + GSK (25 μM); and GSK (25 μM). To understand the influence of IL-4 addition on cytokine production, cells were incubated with the following combination of additives: IL-4 (20 ng/mL); LPS (1 μg/mL); IL-4 (20 ng/mL) + LPS (1 μg/mL); and IL-4 (20 ng/mL) for 24 h followed by LPS (1 μg/mL) for 24 h. Untreated BV2 cells were used as a control. Post-incubation, the supernatants were collected and levels of two pro-inflammatory cytokines, TNF-α and IL-6, were measured using the DuoSet ELISA Assay kit (R&D Systems, MN, USA). ELISA data was analyzed using GraphPad Prism 7.

### RNA-sequencing and proteomics

Data for RNA-sequencing and proteomics in BMDMs were acquired and analyzed as mentioned previously (20).

### Statistical Analysis

One-way ANOVA was used for biological replicates. For each data point, triplicate biological replicates, each with triplicate technical replicates, were performed. Circadian rhythmicity was determined using ECHO analysis software. Statistical analyses were performed using GraphPad Prism 7 or Excel software (Microsoft).

## RESULTS

### Expression of clock genes *Per2* and *Bmal1* in BV2 microglia follow a daily oscillation

*Per2* and *Bmal1* expression and protein levels are representative of the two arms of the circadian transcriptional-translational feedback loop (18), and therefore, were employed as indicators of clock activity. To investigate whether the clock was functional in the BV2 microglial cell line, *Per2* and *Bmal1* gene expression were quantified in naïve (resting) BV2 microglial cells. To assess transcriptional levels of both clock genes, samples were collected starting at 16 h post-serum shock (Hours Post Serum Shock 16 or HPS16), every 2 h for 24 h (**Figure 1a**). Total RNA was isolated from each sample, and *Per2* and *Bmal1* transcript levels were measured using RT-qPCR. The Extended Circadian Harmonic Oscillator (ECHO) application (38) was used to identify oscillations in the transcriptional data and determine whether the gene products oscillated with a circadian period. The ECHO algorithm uses an extended solution of the fixed amplitude oscillator that incorporates the amplitude change coefficient and provides detailed information on the nature of circadian oscillations (38). ECHO analysis indicated that both *Per2* (ECHO period = 16 h, ECHO p-value = 8.31 × 10^-13^) (**Figure 1b**) and *Bmal1* (ECHO period = 16 h, ECHO p-value = 1.11 × 10^-10^) (**Figure 1c**) displayed oscillations at the transcriptional level, although these oscillations are shorter than those considered to be circadian. Both *Per2* and *Bmal1* mRNA oscillations had peaks at around HPS20 and HPS36 post serum shock and a trough around HPS28, paralleling what has been seen in other studies of BV2 microglia (39,40). To further analyze the circadian clock components in BV2 microglia, Western blotting was performed using anti-PER2 and anti-BMAL1 antibodies over circadian time to quantify protein levels (**Figure 1d**). Both PER2 (**Figure 1e**) (ECHO period = 16 h, ECHO p-value = 2.7 × 10^-8^) and BMAL1 (**Figure 1f**) (ECHO period = 16 h, ECHO p-value =1.3 × 10^-4^) displayed significant oscillations according to ECHO with periods and phases that corresponded to what was noted for these genes at the transcript level.

**Figure 1.**
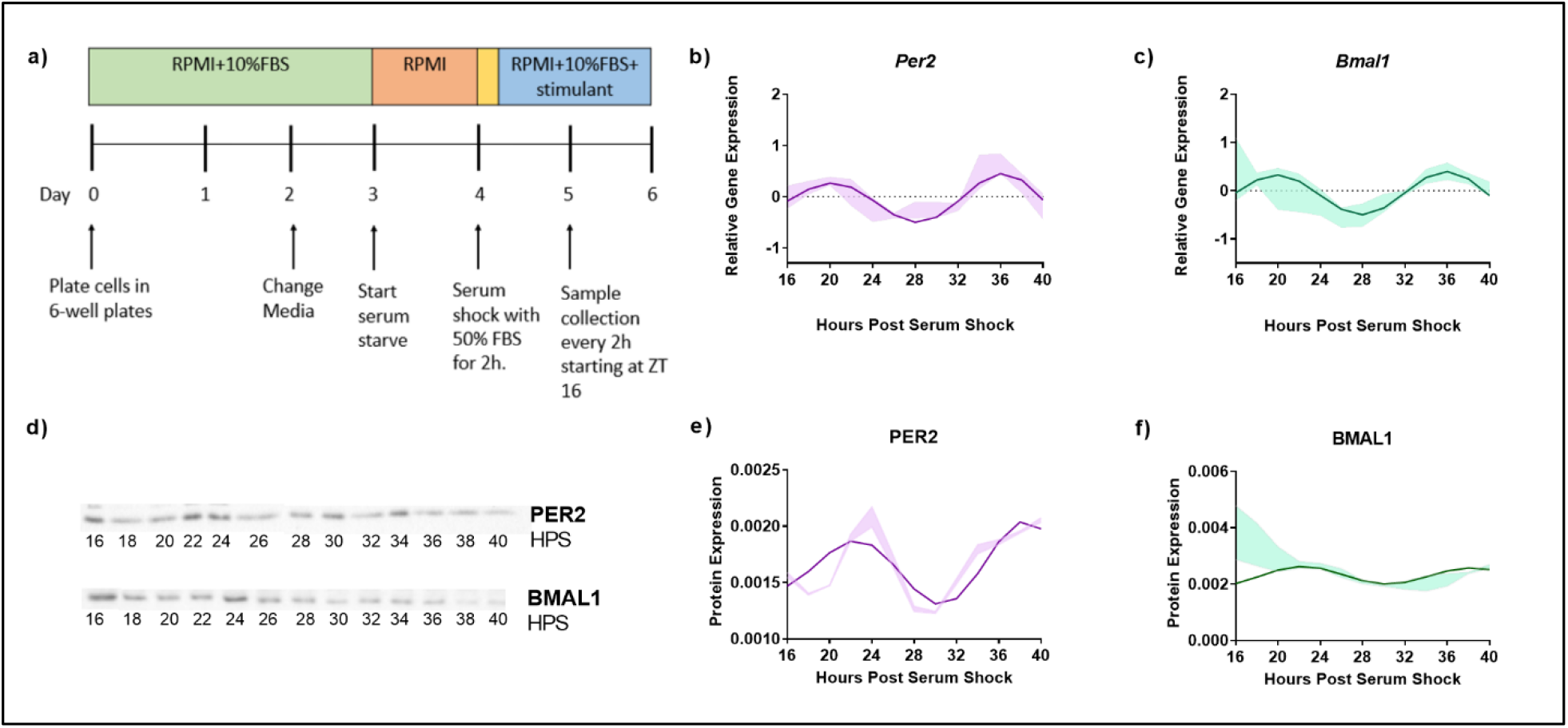
Clock genes *Per2* and *Bmal1* display circadian oscillations in resting BV2 microglia. (a) BV2 microglia were synchronized using the serum-shock protocol and samples were collected every 2 h starting at 16 h post serum-shock (HPS16) for RT-qPCR and Western blotting experiments. (b,c) ECHO fitted plots for mRNA expression of clock genes *Per2* and *Bmal1* in BV2 microglia. Data represented as fold change in expression using Hprt1 as a reference gene and HPS0 as a reference sample for the ΔΔCt method of data analysis (n = 3) (d) Time course samples obtained from serum-shock synchronized BV2 cells analyzed using Western blotting to measure the levels of PER2 and BMAL1. Amido black staining was used to normalize for protein loading. For complete blot images refer Supplementary Information (**Supplementary Figure S3** and **Supplementary Figure S4**) (e,f) ECHO fitted plots for PER2 and BMAL1 protein levels in BV2 cells (n = 3). For all ECHO plots, the bold lines depict the ECHO fitted model and the shaded region represents ±1 standard deviation of model at each time point. Statistical significance was determined using ECHO. All plots had p<0.05 for ECHO significance fit.

### The inflammatory status of the BV2 cell line affects the nature of clock protein oscillations

A functional circadian clock in bone-marrow derived mouse macrophages is receptive to the inflammatory state of the cells (29). Specifically, the activation of bone-marrow derived macrophages into a pro-inflammatory phenotype using LPS, TNF-α, or IFN-γ suppressed the oscillation of core clock components *Per2* and *Nr1d1*, whereas anti-inflammatory activation using IL-4 enhanced their expression (29). To assess whether a similar result is observed in microglia, we subjected BV2 cells to both pro- and anti-inflammatory activation. The bacterial endotoxin LPS is the most commonly used pro-inflammatory stimulus for microglia (41–44). Therefore, to evaluate the influence of LPS exposure on the circadian clock in BV2 microglia, BV2 cells were synchronized using serum-shock, and LPS (1 μg/mL) was added to the growth media post serum-shock. Time course samples for RT-qPCR analysis were collected every 2 h for 24 h starting at HPS16. We found that LPS exposure suppressed the oscillation of the clock genes *Per2* and *Bmal1* in BV2 microglia (**Figure 2a**). This loss of rhythm in the expression of the positive and negative arm components of the circadian feedback loop indicates that the pro-inflammatory activation of BV2 microglia has a suppressive effect on the circadian system, paralleling what was observed in primary macrophages (29).

**Figure 2.**
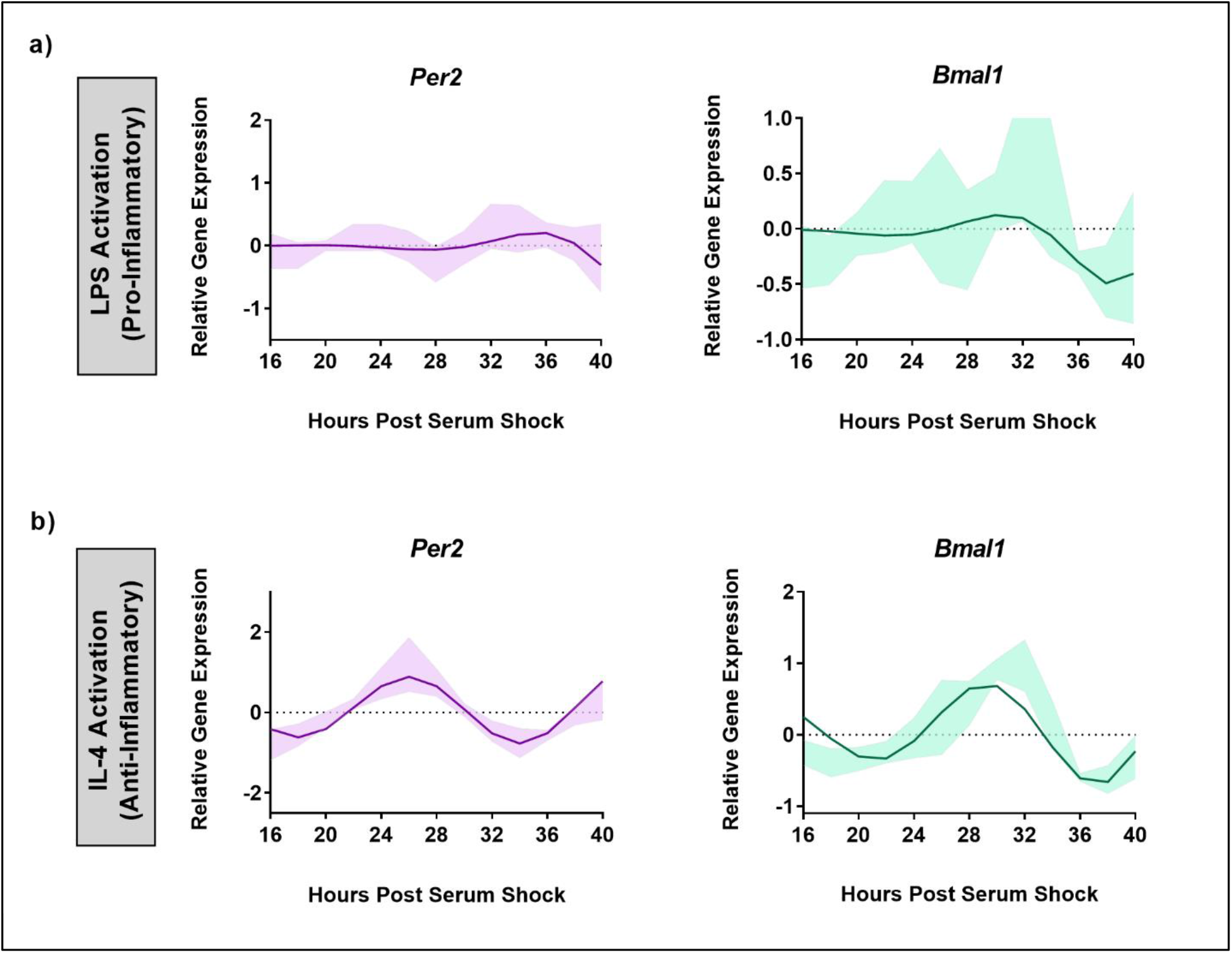
BV2 circadian clock oscillations are affected by the cell’s inflammatory status. ECHO fitted plots for mRNA expression of clock genes *Per2* and *Bmal1* in BV2 microglia (n = 3) under (a) LPS (1 μg/mL) (pro-inflammatory) and (b) IL-4 (20 ng/mL) (anti-inflammatory) activation. Data represented as fold change in expression using Hprt1 as a reference gene and HSP0 as a reference sample for the ΔΔCt method of data analysis. Bold line represent model fit with shaded region representing the standard deviation of model at each time point. All plots had p<0.05 for ECHO significance fit.

The effect of the activation of BV2 microglia into the anti-inflammatory phenotype was assessed by exposing the BV2 cell line to IL-4 (20 ng/mL) after the BV2 cells were synchronized using serum-shock. As with LPS exposure, time course samples for RT-qPCR analysis were collected every 2 h for 24 h starting at HPS16. Unlike for LPS addition, the expression of *Per2* (ECHO period = 16 h, ECHO p-value = 1.04 × 10^−15^) and *Bmal1* (ECHO period = 16 h, ECHO p-value = 2.62 ×. 10^-13^) remained oscillatory (**Figure 2b**) following exposure to IL-4. However, the characteristics of the oscillation of *Per2* and *Bmal1* changed in response to the addition of IL-4. Compared to the resting state, upon IL-4 exposure both *Per2* and *Bmal1* underwent a phase inversion, with a peak around HPS24 to 28 and HPS28 to 32 respectively in the IL-4 activated microglia, as opposed to the peak at around HPS20 for both the genes in the resting state (compare **Figures 1 and 2**). This change in the oscillatory behavior of IL-4 activated BV2 cells vs. resting cells, however, does not abrogate the retention of a daily oscillation in the anti-inflammatory state.

### The BV2 microglial circadian clock times the expression of the NOX2 subunits *p47^phox^* and gp91^*phox*^

A hallmark of a cellular pro-inflammatory state is the presence of oxidative stress, as reflected in the production of ROS (9,45). NOX2 is a major contributor to ROS production leading to oxidative stress (10,14,30,46–49). With the known connection between the clock and inflammation, we hypothesized the circadian clock may control *Nox2* gene expression in BV2 microglia. NOX2 is comprised of six subunits, with gp91^*phox*^ and *p22^phox^* serving as integral membrane proteins that together form the large heterodimeric subunit flavocytochrome b_558_ (cytb_558_) (50), and p40^*phox*^, p47^*phox*^, p67^*phox*^, and Rac together forming the cytosolic subunits (47). Activation of NOX2 occurs though a complex series of protein-protein interactions and translocation (51,52) driven by phosphorylation of p47^*phox*^, which results in attachment of phospho-p47^*phox*^ to the p22^*phox*^ component of cytb558, thus forming an active enzyme complex. For this study, we focused on two essential sub-units of NOX2; p47^phox^ and gp91^phox^ (10).

To assess whether gene expression of *p47^phox^* and *gp91^phox^* was under circadian control in BV2 microglia, time course samples were collected every 2 h for 24 h post-HPS16 from serum-shock synchronized BV2 cells and RT-qPCR was used to measure the expression of *p47^phox^* and *gp91^phox^*. In the resting state, both *p47^phox^* (ECHO period = 16 h, ECHO p-value = 1.7 × 10^11^) and *gp91^phox^* (ECHO period = 16 h, ECHO p-value =1.3 × 10^-17^) displayed significant oscillations, matching the oscillations in *Per2* and *Bmal1* (**Figure 3a**). ECHO plots for both subunits showed peaks around HPS20-24 and HPS36-40, and a trough around HPS28-32, paralleling what was seen for *Per2* and *Bmal1* in the resting state in BV2 cells.

**Figure 3.**
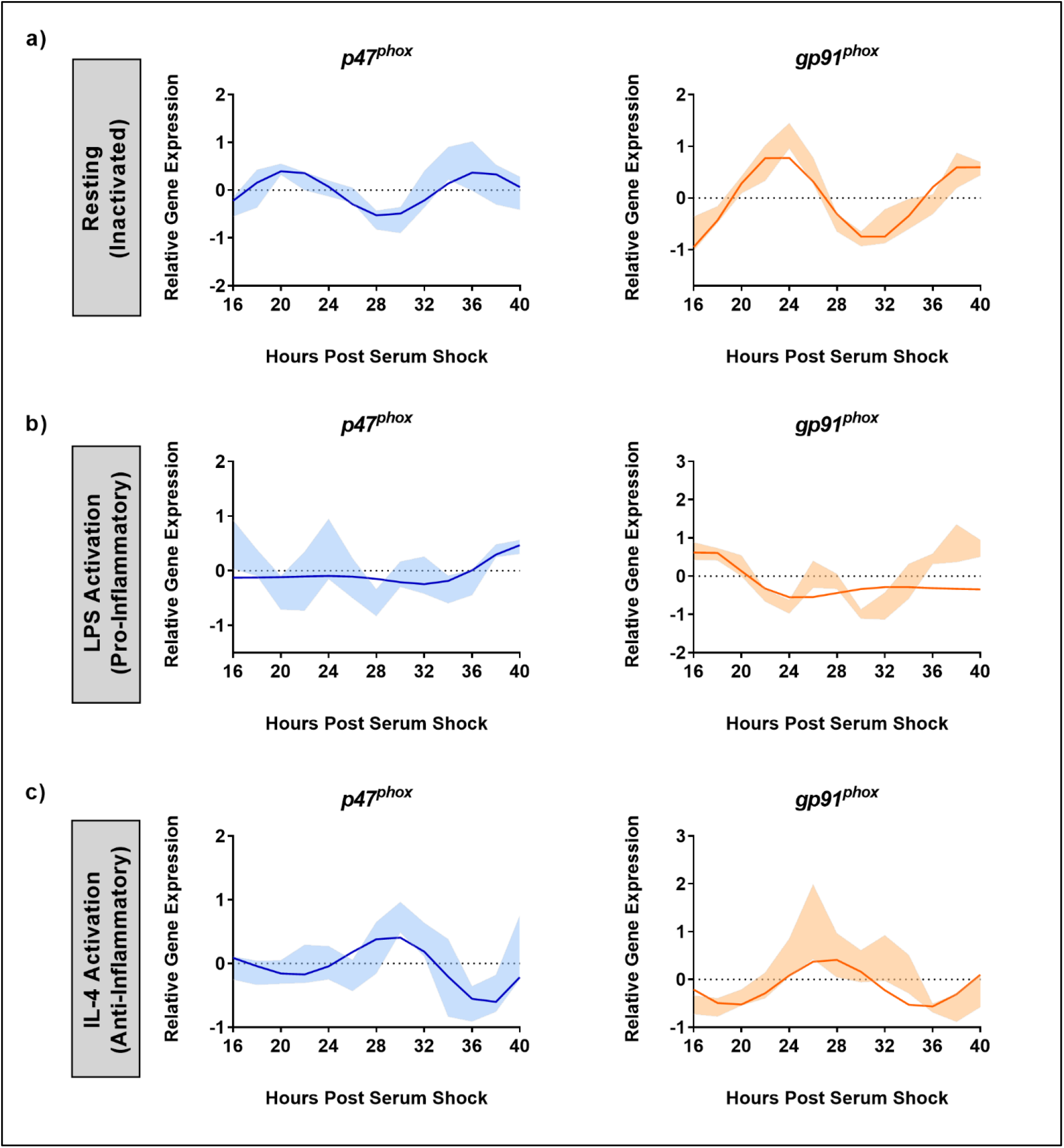
NOX2 sub-units p47^phox^ and gp91^phox^ are under the control of the BV2 microglial circadian clock. ECHO fitted plots for mRNA expression of NOX2 components *gp91^phox^* and *p47^phox^* in BV2 microglia (n = 3) under (a) resting, (b) LPS (1 μg/mL) (pro-inflammatory) and (c) IL-4 (20 ng/mL) (anti-inflammatory) activation. Data represented as fold change in expression using Hprt1 as a reference gene and HSP0 as a reference sample for the ΔΔCt method of data analysis. Bold line represent model fit with shaded region representing the standard deviation of model at each time point. All plots had p<0.05 for ECHO significance fit.

It is known that NOX2 is activated upon addition of LPS (53). Given what we found relating to the effect of LPS on the clock in BV2 microglia, we next assessed the effect of LPS on NOX2 component oscillations. The addition of 1 μg/mL LPS after serum synchronization resulted in the loss of circadian oscillations in both *p47^phox^* and *gp91^phox^* transcript levels (**Figure 3b**). Conversely, the addition of 20 ng/mL IL-4 after serum synchronization did not result in a loss of circadian oscillations for either *p47^phox^* (ECHO period = 16 h, ECHO p-value = 4.98 × 10^-7^) or *gp91^phox^* (ECHO period = 16 h, ECHO p-value = 1.57 × 10^-6^) (**Figure 3c**). Interestingly, there was again a shift in phase between the resting and IL-4 activated states, paralleling what was seen in *Per2* and *Bmal1* after the addition of IL-4. *p47^phox^* expression underwent a phase inversion, with a peak at HPS28-32, as compared to a peak at HPS20 in the resting state. Thus, the two key NOX2 subunits are under circadian regulation in BV2 microglia.

### NOX2 inhibition supports a role for NOX2 in the circadian inflammatory response of BV2 microglia

Given the oscillation of NOX2 and the loss of circadian gene expression both in clock and *Nox2* genes in the pro-inflammatory state, we hypothesized there was a role for NOX2 in clock-driven inflammation. To validate this role for NOX2 in circadian-driven inflammation, we next assessed the effect of NOX2 inhibitors on oscillations in the core clock gene transcripts as well as NOX2 gene expression. To do so, we employed two NOX2 inhibitors to suppress NOX2 function – apocynin (4-hydroxy-3-methoxy-acetophenone) (54–56) and GSK2795039 (GSK, N-(1-Isopropyl-3-(1-methylindolin-6-yl)-1H-pyrrolo[2,3-b]pyridin-4-yl)-1-methyl-1H-pyrazole-3-sulfonamide) (57). Apocynin is known to block the translocation of *p47^phox^* to the membrane and its interaction with membrane-bound p22^*phox*^, thereby preventing the formation of the active enzyme complex (58). Apocynin in not NOX2 specific but is used indiscriminately as a broad NOX inhibitor (59). GSK is a competitive reversible inhibitor of NADPH binding to cytb_558_ *in vitro* and *in vivo* (57), and is selective for NOX2 as compared to the other NOX isoforms, xanthine oxidase, and endothelial nitric oxide synthase (57).

As a proxy for the activity of NOX2 in the presence of LPS and inhibitors, we determined ROS levels via Amplex Red (extracellular) and DCFDA (intracellular) assays. For both assays, additives such as LPS, apocynin or GSK were introduced to serum-shock synchronized BV2 at HPS20 and ROS was measured upon exposure of the BV2 cells to LPS only, LPS with either apocynin or GSK, or no additive (control) for 2 h. In the presence of LPS, both extra- and intracellular ROS levels in BV2 microglia increased relative to the control (**Figures 4a and b**). In the absence of LPS, neither apocynin nor GSK influenced the levels of ROS produced (**Figures 4a and b**). However, in the presence of both LPS and either apocynin or GSK, the levels of both extra- and intracellular ROS were significantly reduced relative to the presence of LPS alone, and were even significantly lower than the control (**Figures 4a and b**). These results suggest that the inhibition of NOX2 may prevent ROS production in BV2 microglia as compared to even basal levels as well as in the presence of LPS.

**Figure 4.**
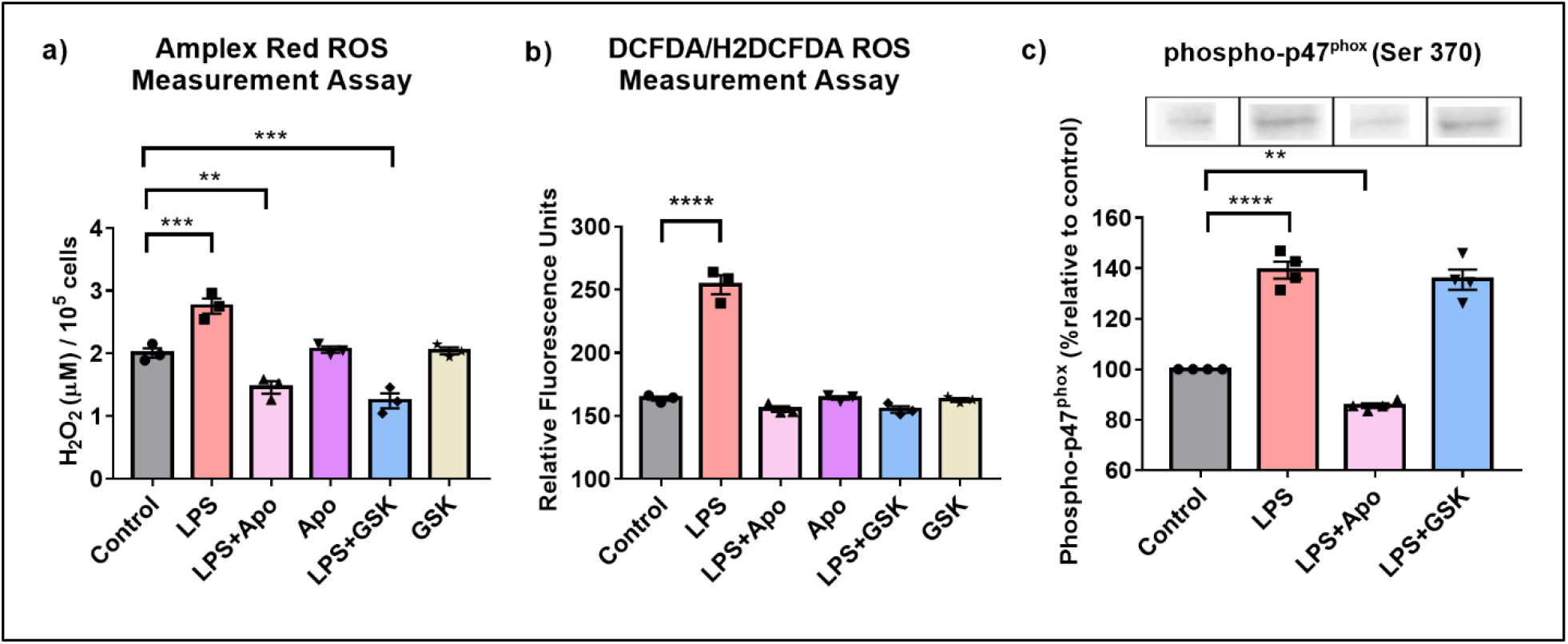
NOX2 inhibition in BV2 microglia results in the reduction of ROS levels and affects phosphorylated-p47^phox^ levels. ROS levels in BV2 microglia with and without LPS and, NOX2 inhibitors apocynin and GSK2795039 determined using (a) Amplex Red Assay to measure extracellular hydrogen peroxide levels and (b) DCFDA Assay to measure intracellular ROS levels. (c) Western blotting analysis of phosphorylated-p47^phox^ levels in BV2 microglia in the presence of LPS and NOX2 inhibitors apocynin and GSK2795039 (n = 3 to 4). For all experiments, BV2 cells with no additives were used as control sample. Data are represented as mean ± SEM and analyzed using single-factor ANOVA test. * denotes p<0.05, ** denotes p<0.01 and **** denotes p<0.0001. For complete blot images, refer Supplementary Information (**Supplementary Figure S5**).

A key step in the activation of NOX2 is the phosphorylation of the p47^*phox*^ subunit (47). To confirm that inhibition of ROS levels by apocynin was NOX2 dependent, even in the presence of LPS, Western blots were obtained to quantify levels of phospho-p47^*phox*^ upon LPS and apocynin addition (**Figure 4c**). In the presence of LPS, there was a 40% increase in phospho-p47^*phox*^ levels. In the presence of LPS + apocynin, however, levels of phospho-p47^*phox*^ fell to approximately 15% below that of the control. These results correlated with extracellular ROS levels (**Figure 4a and 4b**). The reduction in phospho-p47^phox^ levels in the presence of LPS + apocynin suggests that apocynin might regulate ROS production upstream of p47^*phox*^ phosphorylation (60–62). Interestingly, when we repeated the experiment with LPS + GSK, we found that GSK did not affect phospho-p47^*phox*^ levels (**Figure 4c**), which was not unexpected as GSK inhibits the NOX2 cytochrome downstream of the phospho-p47^*phox*^ membrane translocation. Overall, these results confirm effective reduction in ROS levels due to microglial pro-inflammatory activation through the inhibition of NOX2 in the presence of apocynin or GSK, suggesting that NOX2 is the primary source of ROS in BV2 microglia.

Having established that BV2 cells express an active NOX2 that is inhibited by apocynin and GSK, we next studied the effect of NOX2 inhibition on clock gene expression to validate a role for NOX2 in clock-driven inflammation. Serum-shock synchronized BV2 cells were exposed to LPS (1 μg/mL) in the presence of apocynin (100 μM) or GSK (25 μM). Time course samples were collected every 2 h for 24 h starting at HPS16. Addition of apocynin under LPS activation rescued the oscillations in *Per2* (ECHO period = 16 h, ECHO p-value = 1.17 × 10^-6^) and *gp91^phox^* (ECHO period = 16 h, ECHO p-value =1.03 × 10^-12^), which showed oscillations consistent with naïve BV2 microglia (**Supplementary Figure S1a and d**). Conversely, *Bmal1* and *p47^phox^* expression levels no longer showed significant oscillations (**Supplementary Figure S1b and c)**. However, the addition of LPS + GSK rescued oscillations in all four genes: *Per2* (ECHO period = 16 h, ECHO p-value = 2.06 × 10^-12^), *Bmal1* (ECHO period = 16 h, ECHO p-value = 1.02 × 10^−17^), *p47^phox^* (ECHO period = 16 h, ECHO p-value = 5.46 × 10^−5^) and *gp91^phox^* (ECHO period = 16 h, ECHO p-value = 1.48 × 10^-10^), displaying oscillations consistent with naïve BV2 macrophages at the transcriptional level (**Figure 5**). This retention of oscillations under NOX2 inhibition, particularly with GSK, suggests that NOX2 might play a circadian regulatory role in BV2 microglia by facilitating the transition into a pro-inflammatory phenotype.

**Figure 5.**
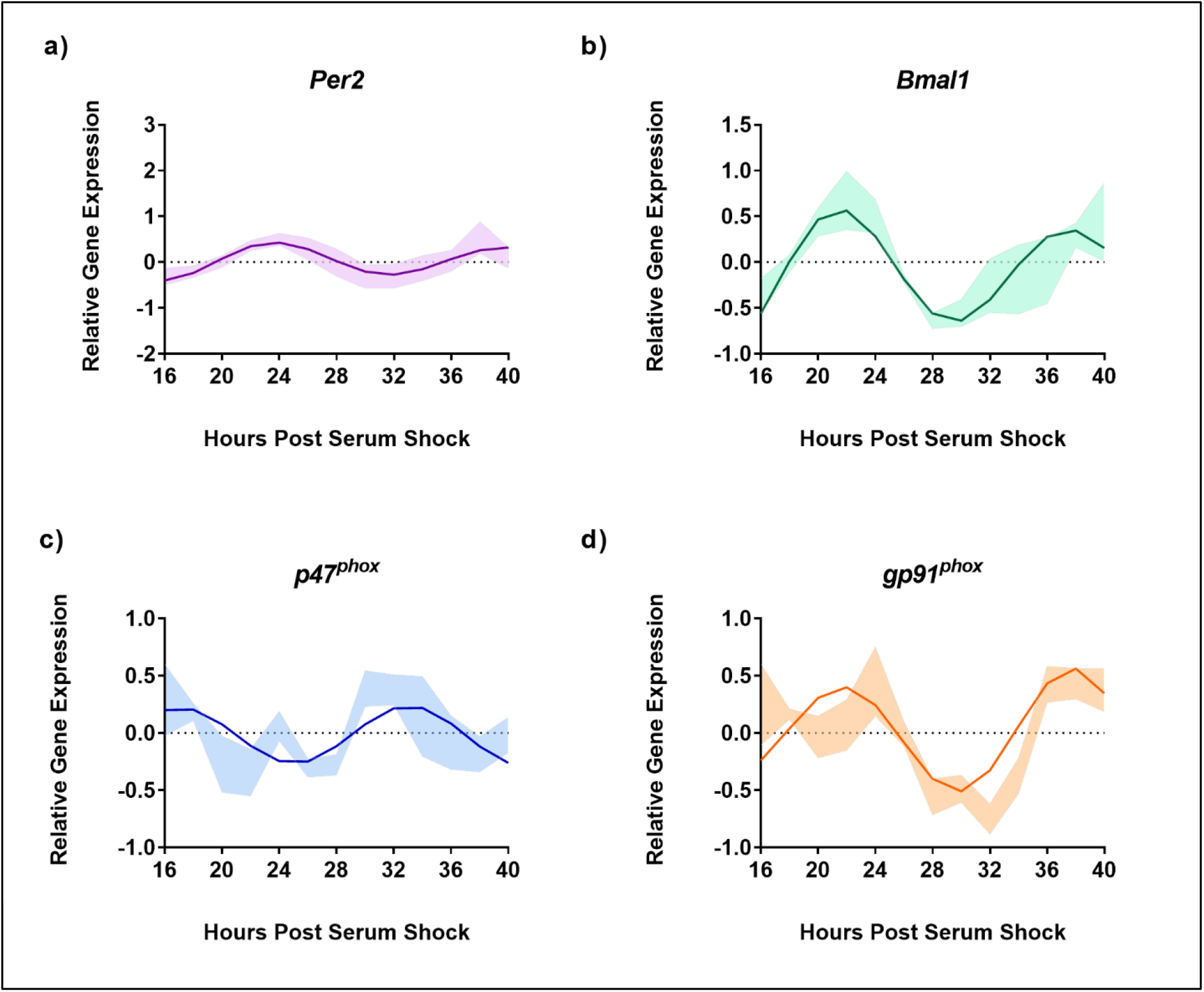
Inhibition of NOX2 by GSK2795039 under LPS activation results in the retention of circadian oscillations in BV2 microglia. ECHO fitted plots for mRNA expression of clock genes (n =3) (a) *Per2* and (b) *Bmal1*, and NOX2 components (c) *gp91^phox^* and (d) *p47^phox^* in BV2 microglia in the presence of 1 μg/mL LPS and 25 μM GSK2895039. Data represented as fold change in expression using Hprt1 as a reference gene and HSP0 as a reference sample for the ΔΔCt method of data analysis. Bold line represent model fit with shaded region representing the standard deviation of model at each time point. All plots had p<0.05 for ECHO significance fit.

### NOX2 regulates the circadianly-induced transition of bone-marrow derived macrophages into a pro-inflammatory state

To validate that the results obtained in BV2 microglia were consistent in primary cells, we next studied the effect of NOX2 inhibition on the circadian clock in bone marrow-derived macrophages (BMDMs). We searched our previous quantitative RNA-sequencing and proteomic analysis datasets (20) to assess the oscillation of *p47^phox^* and *gp91^phox^* in BMDMs. At the transcriptional level, *Per2* (ECHO period = 23.1 h, ECHO p-value = 3.89 × 10^-20^) and *Bmal1* (ECHO period = 29.3 h, ECHO p-value = 3.96 × 10^-20^) showed significant oscillation along with *p47^phox^* (ECHO period = 19.5 h, ECHO p-value = 1.37 × 10^-4^), though *gp91^phox^* had an oscillation beyond the circadian range (ECHO period = 36 h, ECHO p-value = 6.82 × 10^-8^) (**Supplementary Figure 2a**). However, analysis of the proteomic dataset revealed that along with PER2 protein levels (**Figure 6a**), BMAL1 (ECHO period = 28.9 h, ECHO p-value = 1.32 × 10^-5^), and gp91^phox^ (ECHO period = 19.7 h, ECHO p-value = 6.76 × 10^-^5) oscillated with a circadian period, suggesting the post-transcriptional circadian regulation of gp91^phox^, which is common in murine macrophages (**Supplementary Figure 2b**) (20).

**Figure 6.**
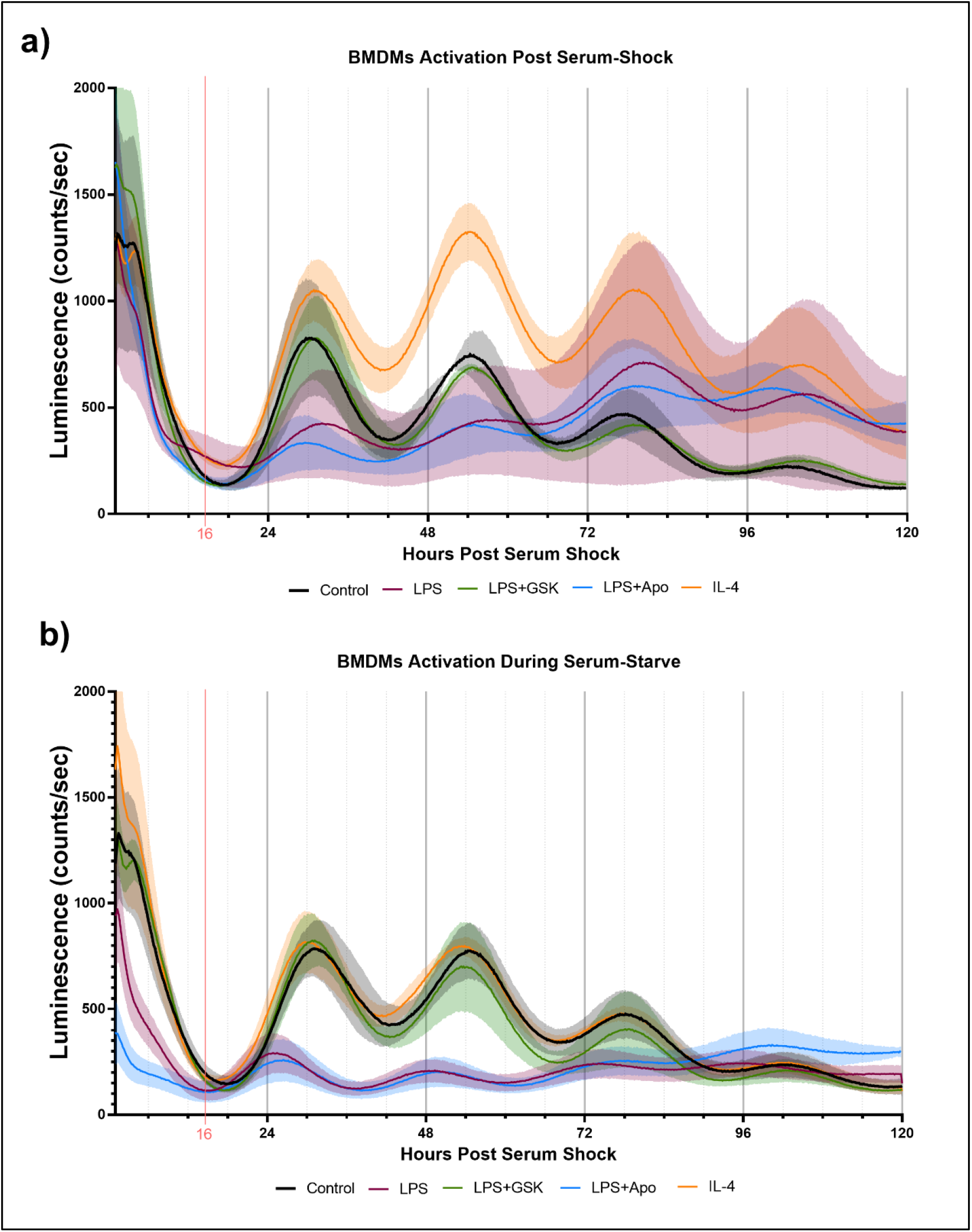
inhibition of NOX2 by GSK2795039 under LPS activation conserves PER2 oscillation in mouse bone marrow-derived macrophages (BMDMs). Luminescence traces obtained luciferase measurement of serum-shock synchronized mouse BMDMs with and without LPS, and NOX2 inhibitors. (a) Cells were exposed to LPS with and without NOX2 inhibitors apocynin and GSK2795039 during the starve stage of the synchronization protocol. (b) Cells were exposed to LPS with and without NOX2 inhibitors apocynin and GSK2795039 post serum-shock stage of the synchronization protocol. Luciferase measurements were obtained by the LumiCycle HPS0 to HPS120 (n = 3). Data are represented as mean ± SD.

To assess the role of NOX2 inhibition on the circadian clock in BMDMs, we used luciferase luminescence measurements to track the levels of PER2 in real time. For this, *Per2::Luc* mice, in which a *Luc* gene is fused in-frame to the 3′ end of the endogenous *mPer2* gene, was used (63). This targeted reporter results in the expression of PER2::LUC bioluminescence similar to endogenous *Per2* expression, rendering it as a real-time reporter of circadian dynamics (63). Bone marrow progenitor cells extracted from *Per2:Luc* C57BL/6J mice were differentiated with recombinant M-CSF into BMDMs and serum-shock synchronized, as per a previously established protocol (20,37). PER2::LUC bioluminescence in BMDMs was assayed over several days using a LumiCycle32 with Leibovitz media containing Luciferin (20,37). Samples treated with neither LPS nor NOX2 inhibitors added were used as controls to confirm that PER2 protein levels oscillated with a circadian period in *Per2:Luc* BMDMs (**Figures 6a and 6b**). We next treated these BMDMs with either LPS (1 μg/mL) or IL-4 (20 ng/mL) post-serum shock, as we did with the BV2 microglia, to elicit pro- and anti-inflammatory activation in the BMDMs respectively. We also analyzed the effect of LPS with NOX2 inhibitors (apocynin (100 μM) and GSK (25 μM)) on PER2 levels in BMDMs. When LPS and/or NOX2 inhibitors were added to the BMDMs post serum-shock synchronization, LPS resulted in a reduction of PER2 amplitude compared to the control sample (**Figure 6a**). When apocynin was added in parallel with LPS, there was a similar decrease in the amplitude of the PER2 oscillation. However, samples in which GSK was added in parallel with LPS maintained the amplitude in PER2 oscillations as compared to untreated BMDMs (**Figure 6a**). Conversely, IL-4 addition resulted in an increase in the amplitude of the PER2 oscillation as compared to the control sample.

Unlike what we had carried out with BV2 microglia, previous work on the response of PER2 oscillations in BMDMs to immunological challenge was performed by adding the treatment prior to serum shock (29). Therefore, to mimic this approach, we repeated our assay of PER2 oscillations in BMDMs by treating the cells with immune insults prior to serum shock synchronization. Like what was seen in BV2 microglia and in post serum shock exposure, exposure to IL-4 did not affect PER2 oscillations as compared to the control sample (**Figure 6b**). When added prior to serum synchronization, LPS addition resulted in the reduction of PER2 levels overall, as well as a reduction in the amplitude of the PER2 oscillation, as has been previously reported (**Figure 6b**) (29). When apocynin was added in combination with LPS prior to serum synchronization, the reduction in PER2 levels and amplitude of oscillation was conserved (**Figure 6b**). However, when GSK was added in combination with LPS prior to serum synchronization, PER2 levels and oscillations were similar to that of the control (**Figure 6b**). The ability of GSK to rescue the levels and amplitude of oscillation of PER2 in BMDMs during LPS activation is consistent with our results in BV2 microglia, and further suggests that NOX2 might play a regulatory role in the circadianly-regulated transition of cells from the monocyte lineage into a pro-inflammatory state.

### Effect of NOX2 on the pro-inflammatory profile of BV2 microglia and BMDMs

To further profile the role of NOX2 in the pro-inflammatory activation of BV2 microglia and BMDMs, we quantified the expression of two common inflammatory cytokines, TNF-α and IL-6, in response to LPS treatment with and without NOX2 inhibitors. Serum-synchronized BV2 cells were stimulated using LPS (1 μg/mL) with and without apocynin (100 μM) or GSK (25 μM) at HPS20. We chose HPS20 as a time point both to allow sufficient time for the cells to recover from the serum-shock, to avoid artifactual gene expression that occurs immediately following serum-shock (35), and to overlap with the peak time in *Per2* oscillation as observed in inactivated BV2 microglia (**Figure 1b**). After 24 h incubation starting at HPS20, the supernatant was collected from BV2 microglia and TNF-α and IL-6 levels were measured using ELISA **(Figure 7a and 7b)**. BV2 microglia treated with LPS demonstrated a significant increase of both TNF-α and IL-6 compared to untreated cells. BV2 microglia treated with only apocynin, or GSK showed no significant change in TNF-α and IL-6 levels compared to untreated cells. However, when apocynin or GSK was added individually in parallel with LPS treatment, there was a significant reduction in TNF-α and IL-6 levels relative to samples with LPS treatment alone, with the greatest reduction in cytokine levels observed in the GSK-treated cells.

**Figure 7.**
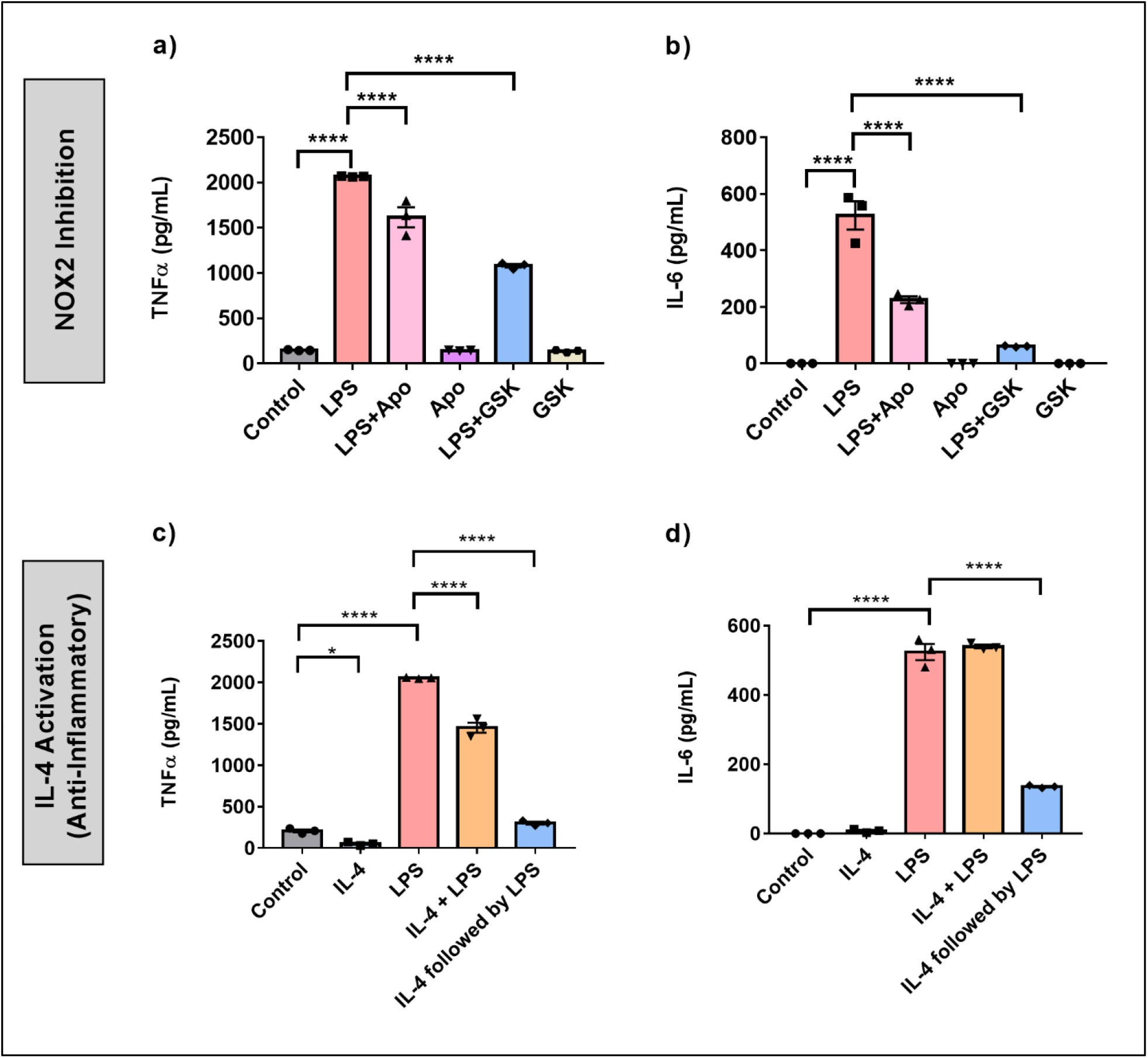
NOX2 inhibition in BV2 microglia creates an inflammatory environment similar to anti-inflammatory activation. (a) TNF-α and (b) IL-6 (pro-inflammatory cytokines) levels in BV2 cells (n = 3) with and without LPS and NOX2 inhibitors apocynin and GSK2795039 measured using ELISA. (c) TNF-α and (d) IL-6 (pro-inflammatory cytokines) levels in BV2 cells (n = 3) under IL-4 activation with and without LPS measured using ELISA. Data are represented as mean ± SEM and analyzed using single-factor ANOVA test. * denotes p<0.05, ** denotes p<0.01, *** denotes p<0.001 and **** denotes p<0.0001.

We next compared NOX2 inhibition to the anti-inflammatory effect on cytokine production in BV2 microglia. To this end, we assessed the effect of IL-4 activation on TNF-α and IL-6 production in BV2 microglia **(Figures 7c and 7d)**. Serum synchronized BV2 cells were incubated with IL-4 (20 ng/mL) for 24 h starting at HPS20. After a 24 hr incubation with IL-4, the media was replaced with fresh media containing LPS (1 μg/mL). In addition, we treated serum synchronized BV2 cells with LPS and IL-4 simultaneously or individually and incubated the cells for 24 h starting at HPS20. While TNF-α levels showed a slight, albeit significant, drop in levels in the presence of IL-4 **(Figure 7c)**, no significant change in IL-6 levels was observed **(Figure 7d)**. Co-treatment of LPS and IL-4 resulted in significant reduction of TNF-α levels compared to samples that had only LPS, IL-6 levels remained unchanged **(Figure 7d)**. Interestingly, LPS activation of BV2 cells post-24 h incubation with IL-4 resulted in a significant drop of both TNF-α and IL-6 **(Figures 7c and 7d)**.

We repeated these experiments in BMDMs to study the effect of NOX2 inhibition on pro-inflammatory activation in primary cells. Serum shock synchronized BMDMs were stimulated using LPS (1 μg/mL) with and without apocynin (100 μM) or GSK (25 μM) at HPS20. After a 24 h incubation, the supernatant was collected and TNF-α and IL-6 levels were measured using ELISA **(Figure 8a and 8b)**. BMDMs treated with LPS demonstrated a significant increase in TNF-α and IL-6 levels compared to the untreated control (**Figures 8a and 8b)**. Concordant with what was seen in BV2 cells, NOX2 inhibition by either apocynin or GSK during LPS treatment resulted in a significant reduction in both TNF-α and IL-6 levels as compared to BMDMs exposed only to LPS. While LPS + GSK showed a more significant drop in TNF-α and IL-6 levels than LPS + apocynin in BV2 microglia, such a difference was not observed in the BMDMs. Compared to BMDMs treated with LPS, addition of IL-4 along with LPS resulted in lower levels of IL-6 (**Figure 8b**). Interestingly, addition of apocynin or GSK along with LPS resulted in the secretion of significant amounts of IL-4 in BMDMs compared to naïve BMDMs or BMDMs exposed only to LPS (**Figure 8c**). Comparison of ELISA results from IL-4 activation (**Figure 7c and 7d)** and NOX2 inhibition **(Figure 7a and 7b)** in BV2 microglia and that of mouse BMDMs (**Figure 8)** suggests that inhibiting NOX2 leads to a microglial response that is similar to that of an anti-inflammatory activation. This is consistent with the inhibition of NOX2 using GSK, wherein the circadian clock retained its oscillation for all genes of interest. These results taken together might indicate that NOX2 inhibition leads to an anti-inflammatory response from both microglia and macrophages, which results in the conservation of circadian oscillations even upon pro-inflammatory activation.

**Figure 8.**
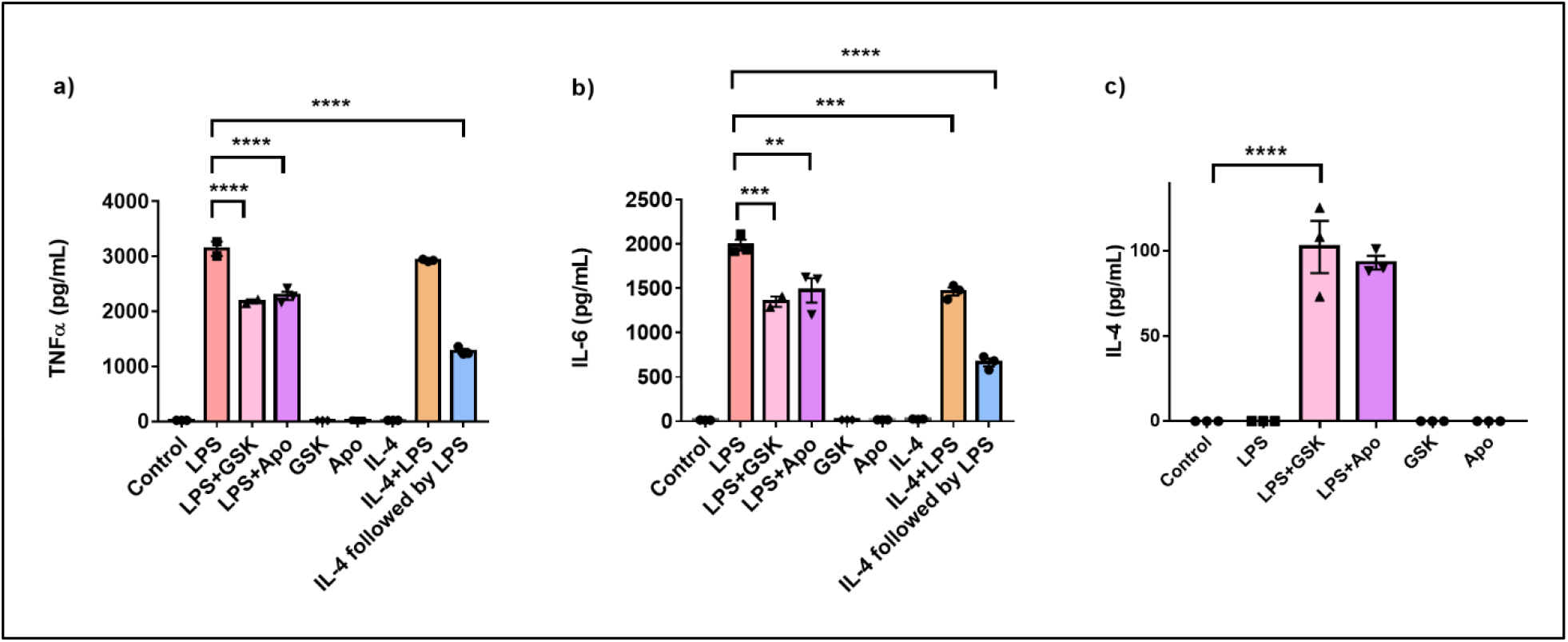
NOX2 inhibition affects the levels of inflammation associated cytokines in mouse bone marrow-derived macrophages. (a) TNF-α and (b) IL-6 (pro-inflammatory cytokines), and (c) IL-4 (anti-inflammatory cytokine) levels in BMDMs (n = 3) with and without LPS and NOX2 inhibitors apocynin and GSK2795039 measured using ELISA. Data are represented as mean ± SEM and analyzed using single-factor ANOVA test. ** denotes p<0.01, *** denotes p<0.001 and **** denotes p<0.0001.

## DISCUSSION

The internal circadian clock comprises a transcription/translation negative feedback loop (TTFL) that controls physiology in macrophages and monocytes to regulate the immune response, playing an important role in the regulation of inflammation, glial activation, oxidative stress, and autophagy (17,22). Concordantly, circadian clock dysfunction has been found to have direct implications on the progression of inflammation, redox defense, and cell death (17). While several studies have been directed towards understanding the impact of circadian regulation on peripheral macrophages, relatively few have focused on microglia. We observed that *Per2* and *Bmal1* gene expression, and PER2 and BMAL1 protein production in serum-shock synchronized BV2 cells displayed circadian oscillations at both transcriptional and protein levels **(Figure 1**). Studies beyond our own have shown that *Per1, Per2, Rev-erb* and *Bmal1* display rhythmic expression, but like our study, have found that while they are predicted to oscillate with a 24 h period in an anti-phasic pattern, this is not the case in the BV2 microglial line (**Figure 1**) (19–21). We predict this is likely due to the circadian modulatory effects commonly noted in cell lines (39,40).

While the effect of the microglial circadian clock on its immune response has been studied (22,40,64–66), the effect of microglial phenotypes on the circadian clock has not been investigated. Pro-inflammatory activation of primary macrophages suppresses *Per2* oscillations, whereas anti-inflammatory activation results in *Per2* amplification (29). We observed similar results for BV2 microglia, wherein the cells lost circadian rhythmicity in *Per2* and *Bmal1* expression upon exposure to LPS (**Figure 2a)** but retained circadian oscillations upon IL-4 exposure **(Figure 2b)**. This suggests that BV2 microglia respond in a similar manner to macrophages and that there is a strong dependence of the microglial clock on the cellular inflammatory state.

Oxidative stress resulting from macrophages or microglial activation is under circadian control (34,67). NOX4 in aortic endothelial cells is influenced by the circadian rhythm (68) and the inhibition of superoxide ameliorated LPS-induced changes in circadian periodicity in peritoneal macrophages (69). NOX2 is associated with the progression of various neurodegenerative diseases (13,16,30,46,70–75). Notably, the inhibition of NOX2 in vascular endothelial cells using diphenyleneiodonium chloride (DPI) restored PER2 rhythmicity under LPS stimulation (76). While these findings linked the circadian clock and oxidative stress, we have shown herein that gene expression of *p47^phox^* and *gp91^phox^* are timed by the clock, indicating that NOX2 is directly timed by the microglial circadian clock. Moreover, we demonstrated that the inflammatory state of the cell has a direct effect on the oscillatory expression of both *p47^phox^* and *gp91^phox^* (**Figure 3**).

While it is clear that NOX2 is controlled by the clock, the reciprocal effect of NOX2 on the clock was unknown. We demonstrated that NOX2 inhibition, by apocynin or GSK, resulted in significant reduction in ROS production and *p47^phox^* phosphorylation when BV2 cells were activated with LPS **(Figure 4)**. Importantly, GSK also rescued rhythmic expression of the two clock genes and two NOX2 genes being tracked **(Figure 5)**. This may indicate that NOX2 plays a crucial role in the transition of microglia from a resting state into a pro-inflammatory phenotype. In parallel, previous studies have shown that *p47^phox^* knockout mice and apocynin-treated mice display reduced levels of cerebrovascular dysfunction and ROS, and increased anti-inflammatory activation of microglia (77). Combined with our data, this suggests that NOX2 may facilitate the link between the circadian recognition of the pro-inflammatory microglial phenotype during neuroinflammation (13).

The results observed in BV2 microglia were further supported by trends obtained in primary mouse BMDMs, with clear oscillations in the levels of *gp91^phox^* and *p47^phox^*. The rescue of PER2 by GSK during LPS treatment directly paralleled what we observed in BV2 microglia (**Figure 6)**. Of note, apocynin failed to improve the amplitude of PER2 oscillations in the presence of LPS. Furthermore, inhibition of NOX2 by apocynin and knockouts of *Nox2* have been shown to reduce pro-inflammatory microglial phenotypes (72). Concordantly, we observed that TNF-α and IL-6 production was significantly reduced in BV2 cells when either apocynin or GSK were present with LPS **(Figure 7a and 7b)**. These results paralleled the reduction in levels of TNF-α and IL-6 as a result of IL-4 priming (i.e., incubated with IL-4 for 24 h) prior to exposure to LPS **(Figure 7c and 7d)**. Similar results were observed when TNF-α and IL-6 levels were measured in BMDMs under NOX2 inhibition, with both cytokines significantly reduced in the presence of GSK along with LPS (**Figure 8a and 8b**). IL-4 levels significantly increased when NOX2 was inhibited using apocynin or GSK in the presence of LPS (**Figure 8c**) indicating that NOX2 inhibition might create a microglial response similar to that of anti-inflammatory activation by IL-4. This could explain why creation of such an anti-inflammatory environment supports retention of the circadian clock in BV2 even in the presence of LPS (**Figure 5 and 6**).

In summary, although neuroinflammation and circadian disruption have been attributed to be major drivers of various neurodegenerative diseases (25), their interrelationship remains unclear. We observed that the BV2 circadian clock depends on the nature of activation of BV2 cells. Furthermore, inhibiting NOX2 resulted in the maintenance of the circadian clock even under pro-inflammatory activation. We suggest that NOX2 is a critical regulator of the microglial inflammatory state and circadian clock function. NOX2 inhibition under pro-inflammatory activation results in the retention of a functional circadian clock in microglia by regulating the inflammatory environment. Our work further suggests a strong link between NOX2 inhibition and the circadian clock in BV2 microglia and primary macrophages that could be further explored in the development of therapeutics to target neuroinflammation.

## ABBREVATIONS

AD: Alzheimer’s Disease
PD: Parkinson’s Disease
Apo: Apocynin
BMDMs: Bone Marrow Derived Macrophages
CT: Circadian Time
DPI: Diphenyleneiodonium chloride
GSK: GSK2795039
HPS: Hours Post Serum Shock
IFN-γ: Interferon Gamma
TNF-α: Tumor Necrosis Factor Alpha
IL-6: Interleukin 6
IL-4: Interleukin 4
NADPH: Nicotinamide Adenine Dinucleotide Phosphate
NOX2: NADPH Oxidase Isoform 2
ROS: Reactive Oxygen Species
TTFL: Transcriptional-Translational Feedback Loop

