## Supplementary Figures for "NOX2 Inhibition Enables Retention of the Circadian Clock in BV2 Microglia and Primary Macrophages"

**SUPPLEMENTARY INFORMATION**

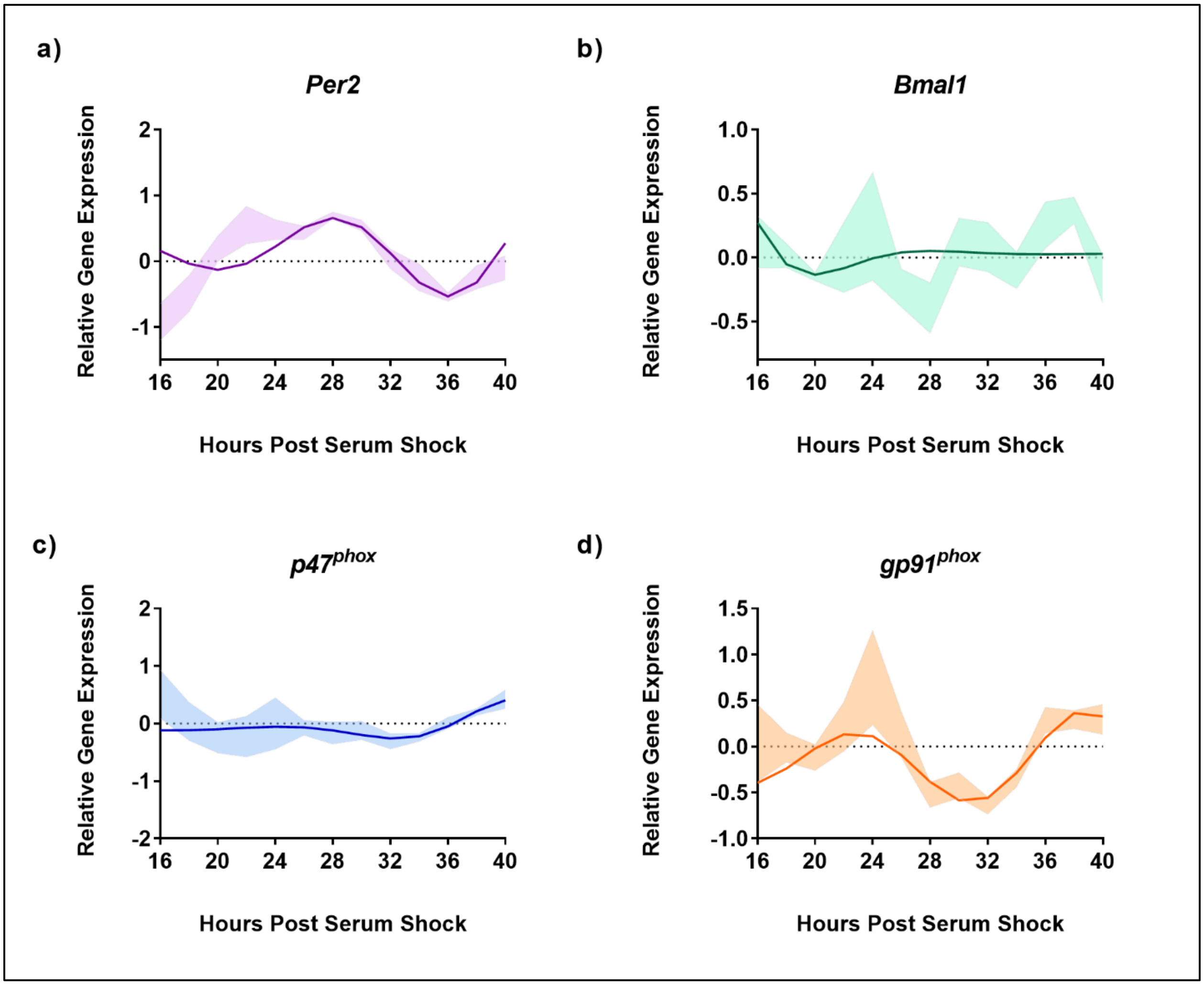


**Supplementary Figure S1. Inhibition of NOX2 by apocynin under LPS activation.** ECHO fitted plots for mRNA expression (n = 3) of clock genes (a) *Per2* and (b) *Bmal1*, and NOX2 components (c) *gp91*$^{phox}$ and (d) *p47*$^{phox}$ in BV2 microglia in the presence of 1 µg/mL LPS and 25 µM GSK2895039. Data represented as fold change in expression using Hprt1 as a reference gene and HPS0 as a reference sample for the ΔΔCt method of data analysis. Bold line represent model fit with shaded region representing the standard deviation of model at each time point. All plots had $p<0.05$ for ECHO significance fit.

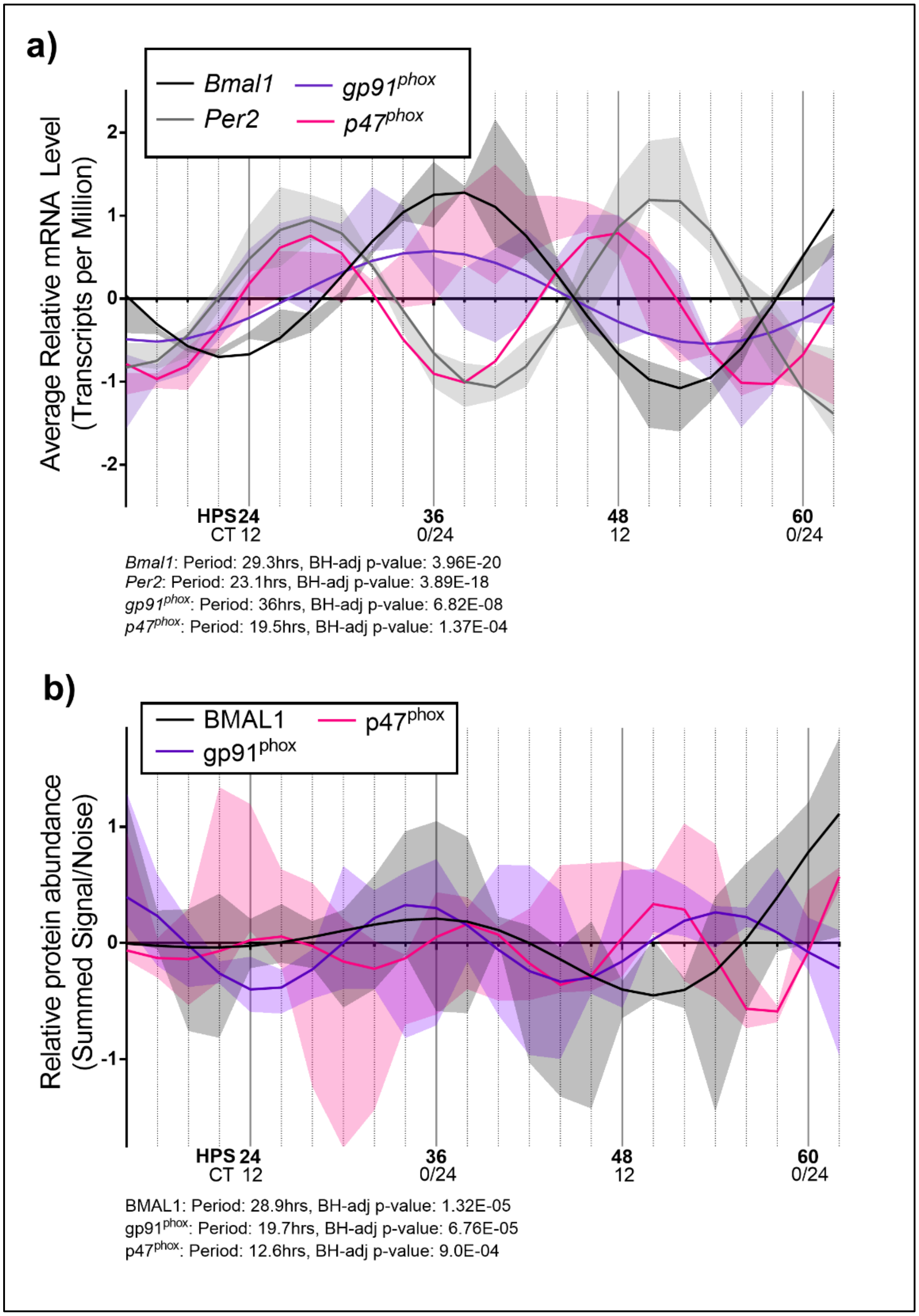


**Supplementary Figure S2. mRNA and protein levels of clock genes and NOX2 components show oscillations in mouse bone marrow-derived macrophages.** Data obtained from previously available RNA sequencing and proteomic analysis datasets showing (a) mRNA oscillations of Per2, Bmal1, $p47^{phox}$ and $gp91^{phox}$ reported in transcripts per million, (b) relative protein abundance oscillations of BMAL1, $p47^{phox}$ and $gp91^{phox}$. Bold line represent model fit with shaded region representing the standard deviation of model at each time point. All plots had $p<0.05$ for ECHO significance fit.

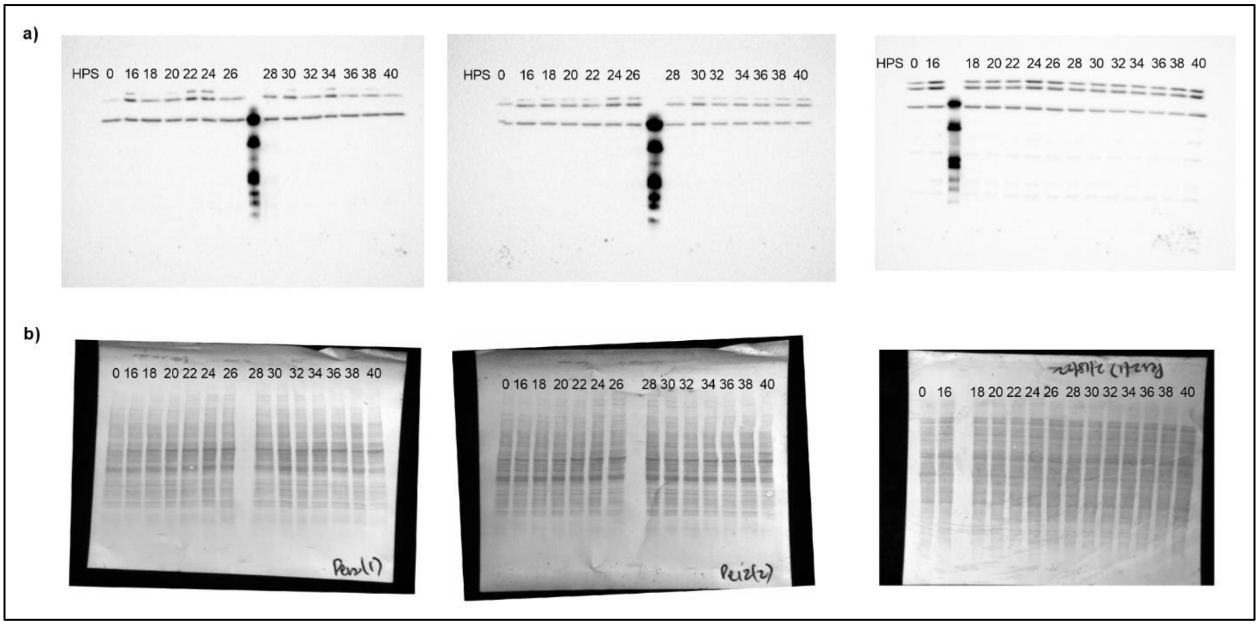


**Supplementary Figure S3. Complete western blot images corresponding to PER2 and its amido black stain in BV2 microglia.** (a) PER2 expression in BV2 microglia for three biological replicates measured every 2 h for 24 h starting at HPS16. (b) Amido black staining of the blots corresponding to PER2 expression in BV2 microglia for three biological replicates measured every 2 h for 24 h starting at HPS16.

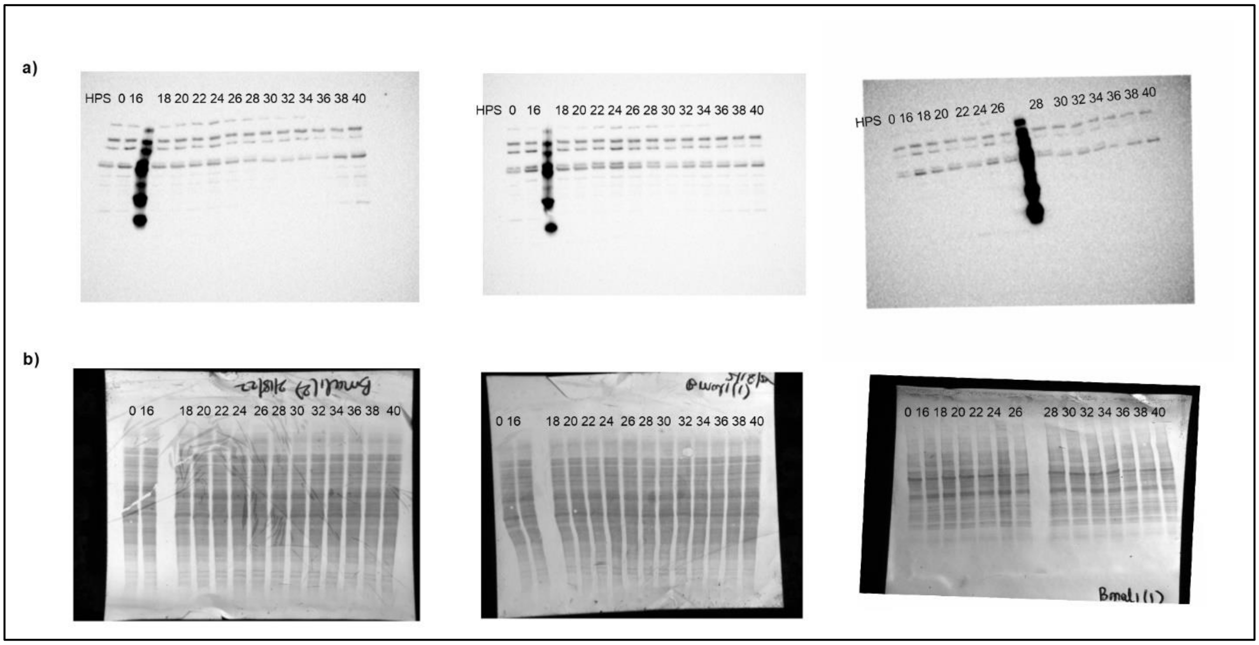


**Supplementary Figure S4. Complete western blot images corresponding to BMAL1 and its amido black stain in BV2 microglia.** (a) BMAL1 expression in BV2 microglia for three biological replicates measured every 2 h for 24 h starting at HPS16. (b) Amido black staining of the blots corresponding to BMAL1 expression in BV2 microglia for three biological replicates measured every 2 h for 24 h starting at HPS16.

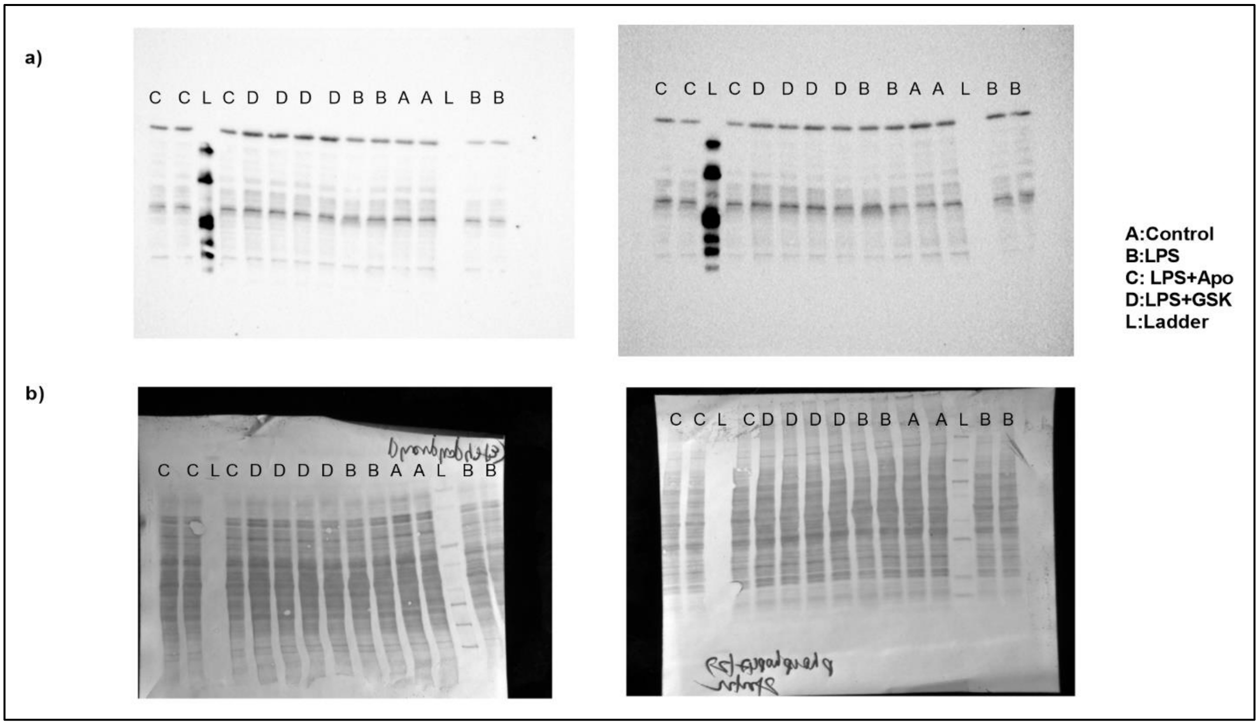


**Supplementary Figure S5. Complete western blot images corresponding to phosphor-p47$^{phox}$(Ser370) and its amido black stain in BV2 microglia.** (a) Phosphorylated-p47$^{phox}$ levels measured in BV2 microglia with and without LPS and NOX2 inhibitors apocynin and GSK2795039, and IL-4 for three biological replicates. (b) Amido black staining of the blots corresponding to Phosphorylated-p47$^{phox}$ levels measured in BV2 microglia with and without LPS and NOX2 inhibitors apocynin and GSK2795039, and IL-4 for three biological replicates.

**Supplementary Table S1. ECHO data for transcript and protein level analysis for clock genes and NOX2 components.** The compiled data including ECHO fitted values and ECHO replicate values for analysis of *Per2, Bmal1, p47$^{phox}$* and *gp91$^{phox}$* genes, and PER2 and BMAL1 protein levels in BV2 microglia.
